# A new law of human perception

**DOI:** 10.1101/091918

**Authors:** Xue-Xin Wei, Alan A. Stocker

**Affiliations:** Department of Psychology University of Pennsylvania

## Abstract

Perception is a subjective experience that depends on the expectations and beliefs of an observer^1^. Psychophysical measures provide an objective yet indirect characterization of this experience by describing the dependency between the physical properties of a stimulus and the corresponding perceptually guided behavior^2^. Two fundamental psychophysical measures characterize an observer’s perception of a stimulus: how well the observer can discriminate the stimulus from similar ones (discrimination threshold) and how strongly the observer’s perceived stimulus value deviates from the true stimulus value (perceptual bias). It has long been thought that these two perceptual characteristics are independent^3^. Here we demonstrate that discrimination threshold and perceptual bias show a surprisingly simple mathematical relation. The relation, which we derived from assumptions of optimal sensory encoding and decoding^4^, is well supported by a wide range of reported psychophysical data^5–16^ including perceptual changes induced by spatial^17,18^ and temporal^19–23^ context, and attention^24^. The large empirical support suggests that the proposed relation represents a new law of human perception. Our results imply that universal rules govern the computational processes underlying human perception.

---

Discrimination threshold is a psychophysical measure that reflects the sensitivity of the observer to small changes in the stimulus variable (Fig. 1A). The threshold depends on the quality with which the stimulus variable is represented in the brain^2^ (*i.e.* encoded - Fig. 1B); a more accurate representation results in a lower discrimination threshold. In contrast, perceptual bias is a measure that reflects the degree to which an observer's perception deviates on average from the true stimulus value (Fig. 1A). Perceptual bias is typically assumed to result from prior beliefs and reward expectations with which the observer interprets the sensory evidence^1^, and thus is determined by factors that are not directly related to the sensory representation of the stimulus. As a result, it has long been believed that there is no reason to expect any lawful relation between perceptual bias and discrimination threshold^3^.

**Figure 1.**
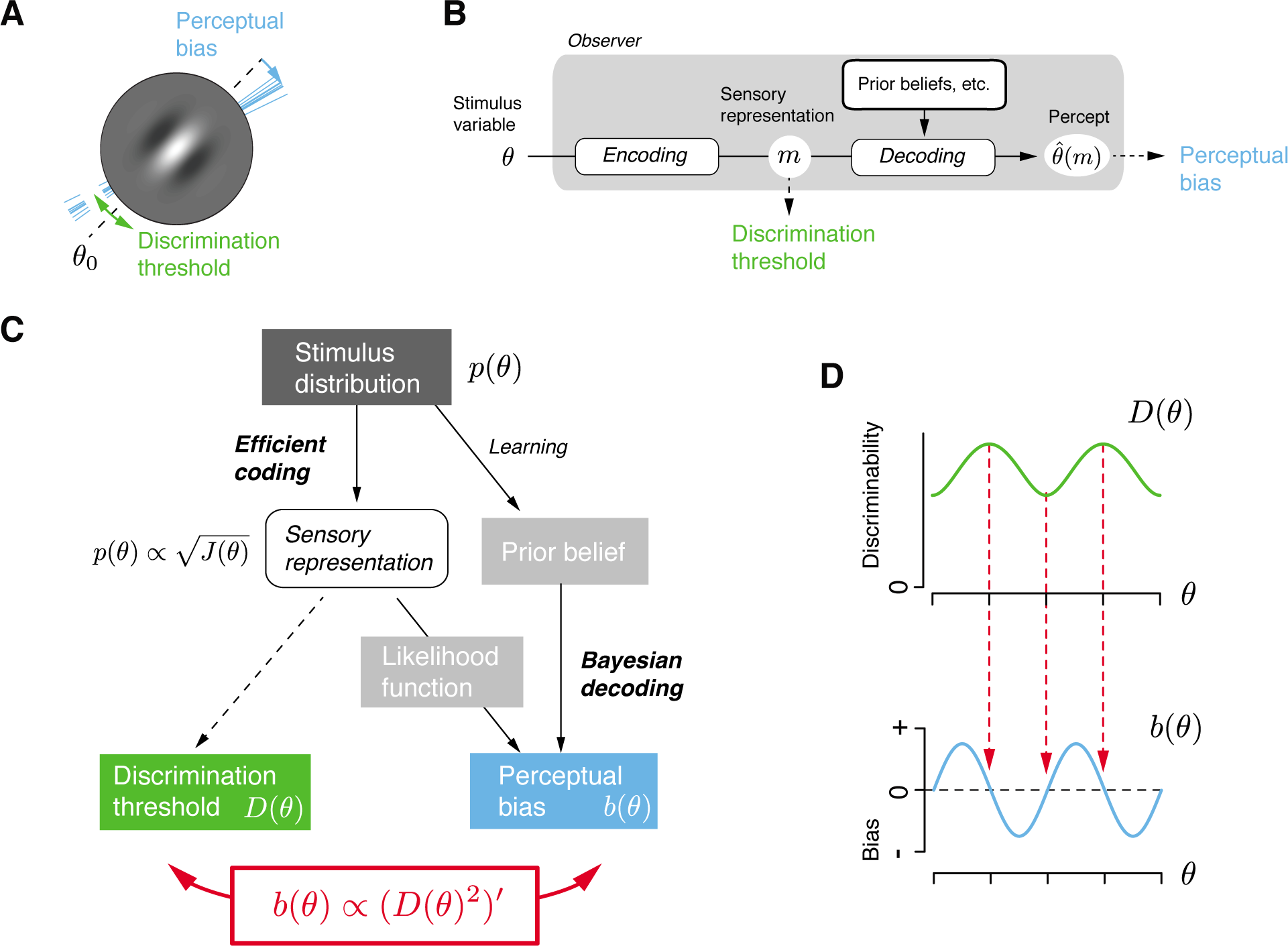
Psychophysical characterization and modeling of perception. (**A**) Perception of a stimulus variable (*e.g.* the perceived local orientation of a visual stimulus) is characterized by discriminability and perceptual bias. Discrimination threshold specifies how well an observer can discriminate small deviations around a particular stimulus orientation *θ*_0_ (green arrows). Perceptual bias specifies how much on average over repeated presentations the perceived orientations (thin blue lines) deviate from the true stimulus orientation (blue arrow). (**B**) Modeling perception as an encoding-decoding processing cascade. Discriminability is limited by the characteristics of the encoding process. Perceptual bias, however, also depends on the decoding process that typically involves cognitive factors such as prior beliefs and reward expectations.(**C**) A new theory of perception proposes that encoding and decoding are optimized for a given stimulus distribution^4^. Within this theory, the encoding accuracy (characterized by Fisher Information *J*(*θ*)) and the bias *b*(*θ*) of the Bayesian decoder are both dependent on the stimulus distribution *p(θ)*. With Fisher information providing a lower bound on discriminability *D*(*θ*)^25^ we can mathematically formulate the relation between perceptual bias and discrimination threshold as *b*(*θ*) « (*D(θ)*^2^)'. (**D**) An arbitrarily chosen, numerical example highlighting the characteristics of the relation: Bias is zero at the extrema of the discrimination threshold (red arrows) and largest for stimulus values where the threshold changes most rapidly.

However, here we derive a simple mathematical relation between discrimination threshold ***D(θ)*** and perceptual bias *b(θ)* based on a recent observer theory of perception^4^. The key idea is that both the encoding as well as the decoding process of the observer are optimally adapted to the statistical structure of the perceptual task (Fig. 1C). Specifically, we assume encoding to be efficient^26^ such that it maximizes the information in the sensory representation about the stimulus given a limit on the overall available coding resources. The assumption implies a sensory representation whose coding resources are allocated according to the stimulus distribution *p(θ).* This is expressed as the encoding constraint 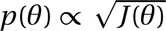 where the Fisher Information *J (θ)* represents the coding accuracy of the sensory representation^4,27–29^. Fisher Information provides a lower bound for the discrimination threshold expressed as 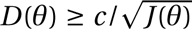 where *c* is a constant^25,30^. Assuming the bound is tight, we can express discrimination threshold in terms of the stimulus distribution as *D*(*θ*) α 1/*p*(*θ*)^28,29^. Furthermore, we have previously shown that the perceptual bias of the Bayesian decoder in the observer model (Fig. 1C) follows 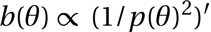, and thus can also be expressed in terms of the stimulus distribution^4,31^. Note that this expression is independent of the details of the loss-function for a large class of symmetric loss-functions (see supplementary Material). Putting all together, we can express a direct relation between perceptual bias and discrimination threshold in the form of

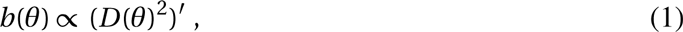

*i.e.* perceptual bias (as a function of the stimulus variable) is proportional to the slope of the discrimination threshold squared. The surprisingly simple mathematical relation predicts that stimulus values with zero bias should correspond to the extrema of the discrimination threshold curve (Fig. 1D, dashed lines). It also predicts that the magnitude of perceptual bias does not necessarily coincide with the magnitude of the discrimination threshold; although intuitively one might have thought that the larger the threshold the larger the noise, and thus the larger the perceptual bias.

We tested the relation against a wide range of existing psychophysical data. Figure 2 shows data for those perceptual variables for which both discrimination threshold and perceptual bias have been reported over a sufficiently large stimulus range. We grouped the examples according to their characteristic bias/threshold patterns. The first group consists of a set of circular variables (Fig. 2A–C). It includes local visual orientation, probably the most well-studied perceptual variable. Orientation perception exhibits the so-called oblique effect^32^, which describes the observation that the discrimination threshold peaks at the oblique orientations yet is lowest for cardinal orientations^5^. Based on the oblique effect, our relation Eq. (1) predicts that perceptual bias at both cardinal and oblique orientations is zero, and that these are the only stimulus values for which the bias is zero. Measured bias functions confirm this prediction^6^. Other circular perceptual variables that exhibit similar patterns are *heading direction* using visual or vestibular information^7^, *2D motion direction* measured with a 2AFC procedure^8,9^ or by smooth pursuit eye-movements^10^, and *motion direction in depth*^11,12^. The relation also holds for the more high-level perceptual variable of perceived *heading direction of approaching biological motion* (human pedestrian)^13^ as shown in Fig. 2C. The second group contains non-circular magnitude variables for which discrimination threshold (approximately) follows Weber's law^33^ and linearly increases with magnitude (Fig. 2D). We predict that these variables should exhibit a perceptual bias that is also (positively or negatively) linear in stimulus magnitude. Indeed, we found this to be true for *spatial frequency* (threshold^5,34^, bias^14^) as well as *temporal frequency* (threshold^35^, bias^16^ - not shown) in vision. Another example is perceived *visual speed* for which discrimination threshold also follows approximately Weber's law^15^ and bias is approximately linear with stimulus speed^16^.

**Figure 2.**
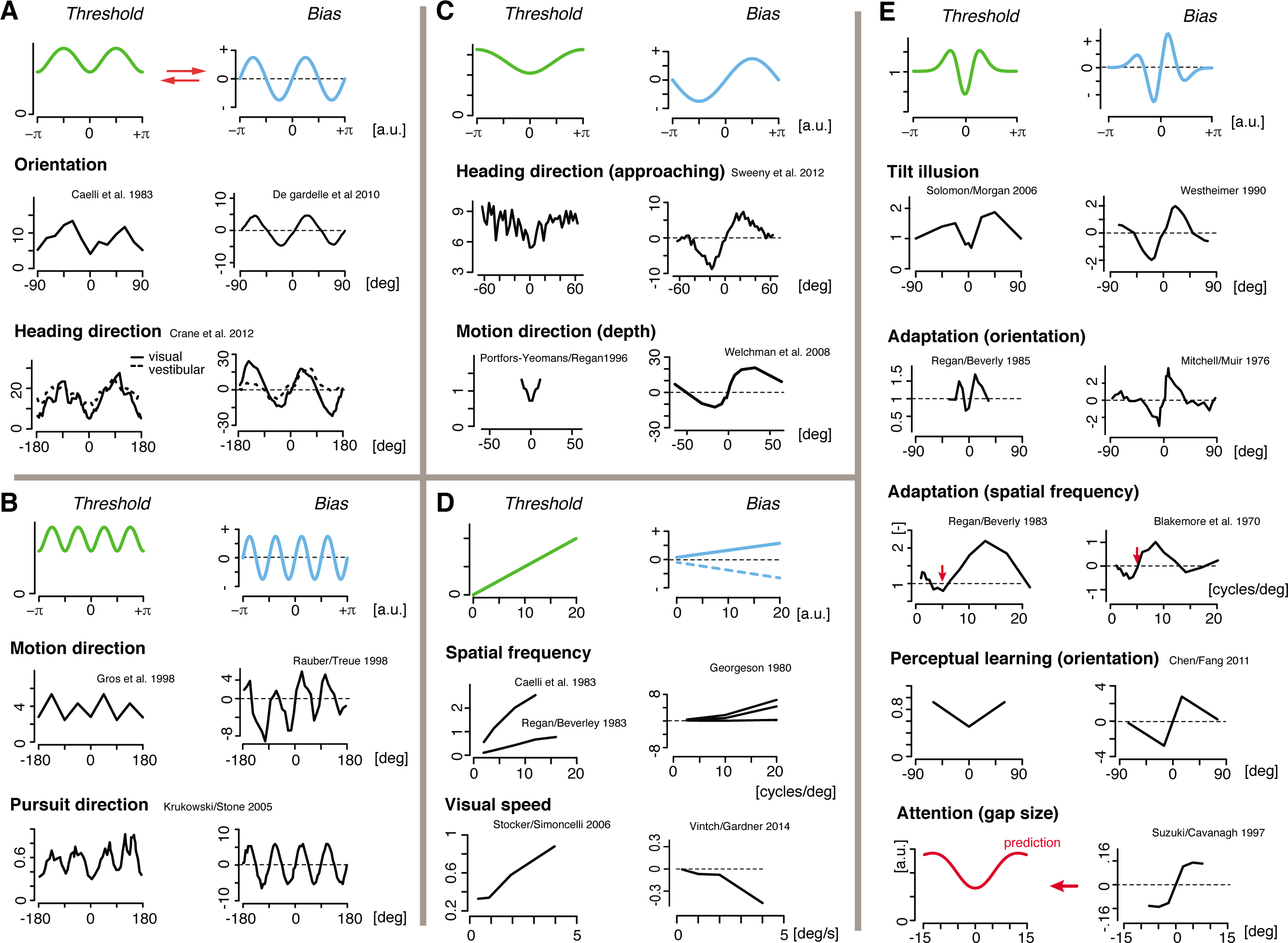
Predicted and measured bias/threshold patterns in perception. Data are organized into different groups. Green/blue curves represent the threshold/bias pattern as predicted by Eq. (1). (**A**) Measured discrimination threshold^5^ and bias^6^ functions for perception of local orientation and heading direction^7^ (solid lines for visual stimulation, dash lines for vestibular stimulation). (**B**) We found similar patterns for perceived motion direction when measured both with a 2AFC procedure^8,9^ or with smooth pursuit behavior^10^. (**C**) The predicted relation also holds for perceived motion direction in depth^11,12^, as well as for the perception of higher-level stimuli such as the approaching heading direction of a person^13^, although the quality of the available data is limited. (**D**) The threshold/bias relation is different for non-circular magnitude variables: discrimination threshold is typically proportional to the stimulus value (Weber’s law), and thus we predict that perceptual bias is also linear in stimulus value. Reported patterns for perceived spatial frequency^5,14^ as well as perceived speed^15,16^ of a visual stimulus match this prediction. (**E**) Bias/threshold patterns resulting from various forms of contextual modulation. Spatial context as in the tilt-illusion^17,18^ and temporal context during adaptation experiments (orientation^19,20^ and spatial frequency^21,22^) lead to similar bias/threshold patterns that are qualitatively well predicted by our theory (red arrows indicate the value of the adaptor stimulus). The predicted relation seems to also hold for perceptual changes induced by perceptual learning^23^ and spatial attention. Spatial attention has been tied to repulsive biases^24^ at the locus of attention and is also known for improvements in discriminability, although little is known how discriminability changes with stimulus value. All data curves are replotted from the corresponding publications except for perceptual bias in orientation^6^ and threshold of visual speed^15^, which both were derived by analyzing the original data.

The last group contains bias/threshold patterns that are not intrinsic to individual specific variables but are induced by contextual modulation (Fig. 2E). Spatial context as in the *tilt-illusion* is known to induce a characteristic repulsive bias pattern in the perceived stimulus orientation away from the orientation of the spatial surround^18^. The corresponding change in discrimination threshold^17^ well matches the predicted pattern based on our theoretically derived relation. Similar bias/threshold patterns have been reported for temporal context, *i.e.,* as *adaptation aftereffects.* Adaptation induced biases and changes in discrimination threshold for perceived visual orientation^19,20^ and spatial frequency^21,22^ nicely match the predicted patterns. At a slightly longer time-scale, *perceptual learning* is also known to reduce discrimination thresholds. We predict that perceptual learning also induces repulsive biases away from the learned stimulus value. This prediction is indeed confirmed by data for learning orientation^23^ and motion direction^36^ (not shown) although the existing data are sparse. Finally, *attention* has been known as a mechanism that can decrease discrimination threshold^37^. We predict that this decrease should coincide with a repulsive bias in the perceived stimulus variable. Although limited in extend, data from a Vernier-gap size estimation experiment support this prediction^24^.

In sum, the derived relation can readily explain a wide array of empirical observations across different perceptual variables, modalities, and contextual modulations. Based on the strong empirical support, we argue that we have identified a new law of human perception. It provides a unified and parsimonious characterization of the relation between discrimination threshold and perceptual bias, which are the two main psychophysical measures that characterize the percept of a stimulus variable.

Only very few *quantitative* laws are known in the perceptual sciences, which include Weber-Fechner’s^2,33^ and Stevens’ law^38^. These laws express simple empirical regularities which provide a compact yet generally valid description of the data. The law we proposed here shares the same virtue. However, unlike these previous laws, the new law is not the result of empirical observations but rather was derived based on theoretical considerations of optimal encoding and decoding^4^. Thus, the law does not merely describe perceptual behavior but rather reflects our understanding of why perception exhibits such characteristics in the first place. The new law allows us to predict either perceptual bias based on measured data for discrimination threshold, or vice versa. This is important because in most cases only one of the two measures has been recorded. One general prediction is that stimulus variables that follow Weber’s law should exhibit perceptual biases that are linearly proportional to the stimulus value as demonstrated with examples in Fig. 2D. Perceptual illusions are often examples of a strong form of perceptual bias. Thus, we predict that these illusions should be accompanied with substantial threshold changes. Perhaps the most surprising result is that the law also holds for contextual modulations (e.g. spatial context) that are instantiated either immediately or on very short time-scales (Fig. 2E). It suggests that changes in encoding and decoding can happen quickly and are matched, which has profound implications for the neural computations and mechanisms underlying perception.

## Acknowledgments

We thank Josh Gold and Michael Eisenstein for helpful comments and suggestions on earlier versions of the manuscript. We would like to express our gratitude to the Office of Naval Research (ONR) for supporting the work (grant N000141110744). We thank V. DeGardelle, S. Kouider and J. Sackur for sharing their data.

## References

1. Hermann von Helmholtz. Handbuch der Physiologischen Optik. Allg. Ezyklopaedie der Physik 9. Bd. Voss, Leipzig, Germany, 1867.

2. Gustav Theodor Fechner. Vorschule der Ästhetik, volume 1. Breitkopf & Härtel, 1876.

3. D.M. Green and J.A. Swets. Signal Detection Theory and Psychophysics. Wiley, New York, 1966.

4. X.-X. Wei and A. A. Stocker. A Bayesian observer model constrained by efficient coding can explain ’anti-Bayesian’ percepts. Nature Neuroscience, 18(10):1509–1517, 2015.

5. Terry Caelli, Hans Brettel, Ingo Rentschler, and Rudi Hilz. Discrimination thresholds in the two-dimensional spatial frequency domain. Vision Research, 23(2):129–133, 1983.

6. Vincent De Gardelle, Sid Kouider, and Jérôme Sackur. An oblique illusion modulated by visibility: non-monotonic sensory integration in orientation processing. Journal of Vision, 10 (10):6, 2010.

7. Benjamin T Crane. Direction specific biases in human visual and vestibular heading perception. PLoS One, 7(12), 2012.

8. Bryan L Gros, Randolph Blake, and Eric Hiris. Anisotropies in visual motion perception: a fresh look. JOSAA, 15(8):2003–2011, 1998.

9. H.-J. Rauber and S. Treue. Reference repulsion when judging the direction of visual motion. Perception, 27:393–402, 1998.

10. Anton E Krukowski and Leland S Stone. Expansion of direction space around the cardinal axes revealed by smooth pursuit eye movements. Neuron, 45(2):315–323, 2005.

11. CV Portfors-Yeomans and D Regan. Cyclopean discrimination thresholds for the direction and speed of motion in depth. Vision Research, 36(20):3265–3279, 1996.

12. Andrew E Welchman, Judith M Lam, and Heinrich H Bülthoff. Bayesian motion estimation accounts for a surprising bias in 3d vision. Proceedings of the National Academy of Sciences, 105(33):12087–12092, 2008.

13. Timothy D Sweeny, Steve Haroz, and David Whitney. Reference repulsion in the categorical perception of biological motion. Vision Research, 64:26–34, 2012.

14. MA Georgeson and KH Ruddock. Spatial frequency analysis in early visual processing [and discussion]. Philosophical Transactions of the Royal Society B: Biological Sciences, 290 (1038):11–22, 1980.

15. Alan A. Stocker and E. P. Simoncelli. Noise characteristics and prior expectations in human visual speed perception. Nature Neuroscience, pages 578–585, April 2006.

16. Brett Vintch and Justin L Gardner. Cortical correlates of human motion perception biases. Journal of Neuroscience, 34(7):2592–604, Feb 2014. doi: 10.1523/JNEUR0SCI.2809-13. 2014.

17. Joshua A Solomon and Michael J Morgan. Stochastic re-calibration: contextual effects on perceived tilt. Proceedings of the Royal Society of London B: Biological Sciences, 273(1601): 2681–2686, 2006.

18. Gerald Westheimer. Simultaneous orientation contrast for lines in the human fovea. Vision Research, 30(11):1913–1921, 1990.

19. D Regan and KI Beverley. Postadaptation orientation discrimination. JOSA A, 2(2):147–155, 1985.

20. Donald E Mitchell and Darwin W Muir. Does the tilt after-effect occur in the oblique meridian? Vision Research, 16(6):609–613, 1976.

21. D Regan and KI Beverley. Spatial-frequency discrimination and detection: comparison of postadaptation thresholds. JOSA, 73(12):1684–1690, 1983.

22. Colin Blakemore, Jacob Nachmias, and Peter Sutton. The perceived spatial frequency shift: evidence for frequency-selective neurones in the human brain. The Journal of Physiology, 210 (3):727–750, 1970.

23. Nihong Chen and Fang Fang. Tilt aftereffect from orientation discrimination learning. Experimental Brain Research, 215(3–4):227–234, 2011.

24. Satoru Suzuki and Patrick Cavanagh. Focused attention distorts visual space: an attentional repulsion effect. Journal of Experimental Psychology: Human Perception and Performance, 23(2):443, 1997.

25. H.S. Seung and H. Sompolinsky. Simple models for reading neuronal population codes. Proceedings of the National Academy of Sciences USA, 90:10749–10753, November 1993.

26. H. B. Barlow. Possible principles underlying the transformation of sensory messages. Sensory Communication, pages 217–234, 1961.

27. N. Brunel and J.-P. Nadal. Mutual information, Fisher information, and population coding. Neural Computation, 10(7):1731–1757, 1998.

28. Deep Ganguli and Eero P Simoncelli. Implicit encoding of prior probabilities in optimal neural populations. In Advances in Neural Information Processing Systems NIPS 23, pages 658–666. MIT Press, 2010.

29. Xue-Xin Wei and Alan A Stocker. Mutual information, fisher information and efficient coding. Neural Computation, 28(2):305–326, February 2016.

30. P. Seriès, A. A. Stocker, and E. P. Simoncelli. Is the homunculus ‘aware’ of sensory adaptation? Neural Computation, 21:3271–3304, December 2009.

31. X.-X. Wei and Alan A. Stocker. Efficient coding provides a direct link between prior and likelihood in perceptual bayesian inference. In Advances in Neural Information Processing Systems NIPS 25, pages 1313–1321. MIT Press, December 2012.

32. S. Appelle. Perception and discrimination as function of stimulus orientation. Psychological Bulletin, 78:266–278, 1972.

33. Ernst Heinrich Weber. The sense of touch. Academic Press, 1978.

34. Fergus W Campbell, Jacob Nachmias, and John Jukes. Spatial-frequency discrimination in human vision. JOSA, 60(4):555–559, 1970.

35. M.B. Mandler. Temporal frequency discrimination above threshold. Vision Research, 24(12): 1873–1880, 1984.

36. S. Szpiro, M. Spering, and M. Carrasco. Perceptual learning modifies untrained pursuit eye movements. Journal of Vision, 14(8):8, 2014.

37. Marisa Carrasco and Brian McElree. Covert attention accelerates the rate of visual information processing. Proceedings of the National Academy of Sciences, 98(9):5363–5367, 2001. doi: 10.1073/pnas.081074098.

38. Stanley S Stevens. On the psychophysical law. Psychological Review, 64(3):153, 1957.

